# Rotating a petavoxel reconstruction exposes the viewing-angle bias inherent to Golgi-Cox and confocal dendritic-spine classification

**DOI:** 10.64898/2026.06.15.732500

**Authors:** Elias Manjarrez, Sandra T. Hernández, Miguel A. Zamora-Ursulo, Amira Flores

## Abstract

Dendritic spines are the principal postsynaptic sites of excitatory transmission. For over a century, their shape has been sorted into discrete categories such as filopodia, thin, long thin, stubby, mushroom, and branched, largely by Golgi-Cox impregnation and, more recently, confocal microscopy. However, both approaches share a fundamental limitation. The histological sectioning and single-viewpoint imaging that these methods rely on cannot control the orientation of a spine relative to the observer. Because a spine is a three-dimensional object, the projection seen depends on how its parent dendrite lies within the section. Here, using the publicly available H01 petavoxel reconstruction of human temporal cortex imaged by serial-section electron microscopy (EM), we show that spine-shape classification depends strongly on viewing angle. A total of 445 spines on layer 4 basal dendrites of five pyramidal neurons were classified from an initial viewpoint (Angle 1), then reclassified after rotation in Neuroglancer (Angle 2). Only 20.9% kept their category, so chance-corrected agreement was negligible (Cohen’s kappa = 0.027). These observations provide direct evidence that the rigid Golgi-Cox and confocal taxonomies conflate true spine morphology with the arbitrary angle of view. Our results, therefore, support recasting spine shape as a three-dimensional continuum, measurable in petavoxel reconstructions such as H01 through free rotation in Neuroglancer.

**Significance statement:** The classification of dendritic spines into discrete shape classes underpins a vast literature on synaptic plasticity, development, and disease. Yet it rests on two-dimensional images whose viewing angle is not controlled. By rotating the same human spines in a nanoscale EM reconstruction, this study shows that four out of five spines change category with viewpoint alone. The finding exposes a systematic bias in Golgi-Cox and confocal classifications. It argues that spine morphology should be treated as a measurable three-dimensional continuum rather than a set of fixed labels.

## Introduction

Dendritic spines are small, membranous protrusions that receive the great majority of excitatory synaptic inputs in the mammalian cortex. Their size and shape are tightly coupled to synaptic strength and plasticity (Arellano et al., 2007; Rochefort and Konnerth, 2012). Since Cajal first drew them with the Golgi method (Yuste, 2015), spines have conventionally been grouped into a small number of morphological classes (Peters and Kaiserman-Abramof, 1970). These are typically filopodia, thin, long thin, stubby, mushroom, and branched, and the grouping assumes that each class reflects a distinct functional state (Arellano et al., 2007; Bourne and Harris 2008; Rodriguez et al. 2008; Garcia-Lopez et al., 2010; Risher et al., 2014; Berry and Nedivi, 2017; Tendilla-Beltran et al., 2019; Otero et al., 2022; Ustinova et al., 2024; Flores et al., 2025; Hu et al., 2025; Aguilar-Hernandez et al., 2025; Paul et al., 2025; Masoli et al., 2025; Doliwa et al., 2025; Escribano-Cadena et al., 2025; Kayedi-Bakhtiari and Vatanparast, 2025; Cabrera-Pedraza et al., 2026; Dai et al., 2026; Tennin et al., 2026). This taxonomy remains the working language of the field, and it is built almost entirely on Golgi-Cox impregnation or on confocal microscopy of fluorescently labeled neurons.

However, both dominant techniques share a decisive limitation. They reconstruct a three-dimensional object from images whose orientation cannot be controlled. In a Golgi-Cox preparation, the plane of histological section is fixed before the tissue is imaged. A spine is therefore observed from whatever angle its parent dendrite happens to adopt in the slice. A filopodia spine that projects toward or away from the observer can collapse into the silhouette of a stubby protrusion, and two crossing spines can appear as a single branched structure. Confocal and two-photon microscopy impose a further constraint of poor axial resolution. The z dimension, which is precisely the one most often hidden from the observer, is therefore the least reliable (Pchitskaya and Bezprozvanny, 2020). Classification criteria such as head width, neck length, and the length-to-width ratio are all measured in the image plane. The assigned class is therefore a property of the projection rather than of the spine itself.

A growing body of quantitative work has independently questioned whether discrete spine classes exist at all, and the view that spine morphology forms a continuous spectrum rather than a set of distinct categories has become increasingly accepted. Analyses of three-dimensional reconstructions by confocal microscopy in human and mouse cortex (Ofer et al. 2022) and of spine necks by EM (Ofer et al. 2021) consistently report unimodal, continuous distributions of head volume, neck length, and neck diameter, with no evidence of separable subtypes. This continuum is further supported by different techniques, which find spine shapes distributed along a gradient rather than falling into discrete classes (Wilson et al., 1983; Portera-Cailliau et al., 2003; Benavides-Piccione et al., 2012; Tonnensen et al., 2014; Arellano et al., 2007; Bhatt et al., 2009; Loewenstein et al., 2011; Reberger et al., 2018; Ofer et al., 2021; 2022). To capture such diversity without imposing fixed labels, a variety of clustering algorithms and criteria for selecting the optimal number of clusters have been explored, although which approach is most suitable remains unsettled (Bokota et al. 2016; Luengo-Sanchez et al. 2018; Urban et al., 2019; Kashiwagi et al., 2019; Pchitskaya and Bezprozvanny 2020; Pchitskaya et al. 2023; Choi et al., 2023; Ferreira et al. 2024; Reimer et al., 2024; Gilles et al., 2024; Theobald et al., 2025; Smirnova et al., 2026).

Reviews of the segmentation and classification literature reach the same conclusion and advocate replacing rigid classes with clustering of continuous descriptors (Pchitskaya and Bezprozvanny, 2020; Pchitskaya et al., 2023). One thing has been missing, however. No study has shown directly that the act of classification itself is unstable. No study has shown that the same spine, measured by the same observer, can be assigned to different classes solely based on the viewing angle of its dendrite. Such a demonstration requires a substrate on which a single spine and its dendrite can be inspected, rotated freely, and reclassified. Neither fixed Golgi-Cox sections nor conventional confocal stacks can provide this.

The recent reconstruction of a cubic millimeter of human temporal cortex using serial-section EM, known as the H01 dataset (Shapson-Coe et al., 2024), now makes this possible. H01 comprises 1.4 petabytes of data, about 57,000 cells, and about 150 million synapses. It is distributed through the browser-based Neuroglancer viewer, which permits arbitrary three-dimensional rotation of segmented dendrites at nanometer resolution. This resource, therefore, allows a spine to be classified from one viewpoint and then classified again after rotation of its dendrite. The contribution of viewing angle to the classification outcome can thus be isolated while all other variables are held constant.

The present study uses this capability to test a relevant question. If the conventional spine taxonomy captured an intrinsic property of spines, the classification should be invariant under rotation. It was therefore hypothesized that rotational, nanometer-scale analysis of human cortical spines positioned in their dendrites would reveal substantial and systematic reclassification as a function of viewing angle. Such a result would expose a bias inherent to Golgi-Cox and confocal classification. It would also support the view that spine morphology is a rotation-invariant continuum rather than a set of discrete categories.

## Materials and Methods

### Dataset

Analyses were performed on the publicly available EM H01 reconstruction of the human cerebral cortex (Shapson-Coe et al., 2024). The sample is a fragment of the anterior middle temporal gyrus, approximately 1 mm^3^, that was resected during surgery to access an underlying epileptic focus. The tissue was rapidly fixed, stained with heavy metals, sectioned at about 33 nm, and imaged by multibeam scanning EM at 4 × 4 nm pixels. The resulting volume was aligned and segmented using flood-filling networks to yield three-dimensional meshes of essentially every neuron and process, and conservative c3 agglomeration was applied.

### Spine visualization and rotation in Neuroglancer

A total of 445 spines on layer 4 basal dendrites of 5 pyramidal neurons were classified by a trained observer from an initial viewpoint (H01 fragment ID: 3808204067, 4096340441, 4460415817, 29223100509, 29994996547). Segmented dendrites were explored in Neuroglancer (developed by J. Maitin-Shepard and the Google Connectomics team), the browser-based viewer distributed with H01. Neuroglancer presents three linked panels. The first is a two-dimensional cross-sectional panel in which any pair of the X, Y, and Z axes can be displayed, and any voxel can be selected. The second is a three-dimensional rendering panel in which the selected object can be freely rotated, panned, and zoomed about a central set of axis guides. The third is a control panel in which cortical layer, cell type, and individual segments are selected, and in which each object’s unique identifier (ID) is displayed, starred, and recovered for later analysis. Dendrites of layer 4 (L4) neurons were retrieved by their segment IDs, and their spine-bearing segments were rendered as three-dimensional meshes.

For each analyzed dendrite, the rendering was first oriented to the position where the segment initially loaded. This is referred to as Angle 1, the original designation, and it corresponds to the kind of single, uncontrolled viewpoint that a Golgi-Cox or confocal preparation imposes. Here, it is important to note that we rotated the dendrite containing spines, not the spines themselves. This differs from previous EM studies, in which the rotation is applied to single spines, only to quantify their geometric shape (Arellano et al., 2007; Ofer et al., 2021). In this context, our study is completely original when compared with previous EM analysis of spines.

Each visible spine included in the dendrite was classified by one trained observer into one of six conventional categories, namely filopodia, long thin, thin, stubby, mushroom, or branched, using standard head and neck criteria (Risher et al., 2014). The same dendrite was then rotated in three dimensions within Neuroglancer to a second orientation, referred to as Angle 2, and each spine was independently reclassified using identical criteria. Because the object, the observer, and the criteria were all held constant, any change in assigned class between Angle 1 and Angle 2 is attributable to viewing angle alone.

It is important to note that, to emulate the uncontrolled viewing geometry of conventional preparations, the two orientations were chosen without reference to the spine shape. In a Golgi-Cox or confocal preparation, the angle at which a dendrite and its spines are presented to the observer is set by the arbitrary plane of histological sectioning, a plane fixed before the spines are seen and unrelated to the morphology of any individual spine.

We reproduced this blindness in two ways. First, Angle 1 was simply the orientation in which each segmented dendrite was initially loaded in Neuroglancer, an arbitrary starting pose determined by the dataset rather than by the trained observer. Second, the rotation to Angle 2 was applied to the dendrite before its individual spines were inspected or classified, so that the second viewing angle was likewise selected independently of the shape of any spine. The trained observer, therefore, did not rotate spines toward a preferred or expected appearance, exactly as a histologist cannot orient an individual spine within a tissue slice. Because the two views were chosen blind to spine morphology, no fixed set of rotation angles was prescribed, and the reclassification reported here reflects the ordinary ambiguity of single viewpoint imaging rather than any chosen geometry.

### Statistical analysis of the confusion matrix

Paired Angle 1 and Angle 2 classifications were tabulated as a 6 × 6 confusion matrix in which rows index the original Angle 1 designation and columns index the post-rotation Angle 2 designation. Diagonal entries represent concordant classifications, where the spine retained its category after rotation. Off-diagonal entries represent discordant classifications, where the spine changed category. Overall concordance was computed as the proportion of classification events falling on the diagonal, that is, the sum of diagonal counts divided by the grand total.

Agreement was additionally corrected for chance using Cohen’s kappa (Cohen, 1960; Sim and Wright, 2005), which adjusts the observed agreement to account for the agreement expected by chance. The directionality of the disagreements was tested using the McNemar-Bowker test of marginal symmetry, a generalization of McNemar’s test (Bowker, 1948), which evaluates whether the off-diagonal pair counts (cell ij versus cell ji) differ more than expected by chance across the table. A McNemar-Bowker statistic that rejects symmetry indicates that reclassification is biased in particular directions, for example, that stubby and mushroom transitions are asymmetric, rather than reflecting random labeling noise. Analyses were performed in MATLAB (The MathWorks, Natick, MA), and p-values below 0.05 were considered significant.

## Results

### Qualitative observations

First, we present representative examples of dendrites before and after rotation, along with their corresponding per-spine labels. Figure 1 traces the path from the human temporal cortex to the three-dimensional rotation of single spines. A cubic millimeter fragment of the anterior middle temporal gyrus (Figure 1a, b) was sectioned with an Automated Tape-Collecting Ultramicrotome (ATUM) and imaged by multibeam scanning electron microscopy (mSEM) (Figure 1c), then aligned and segmented to yield the H01 volume explored in Neuroglancer (Figure 1d).

**Figure 1.**
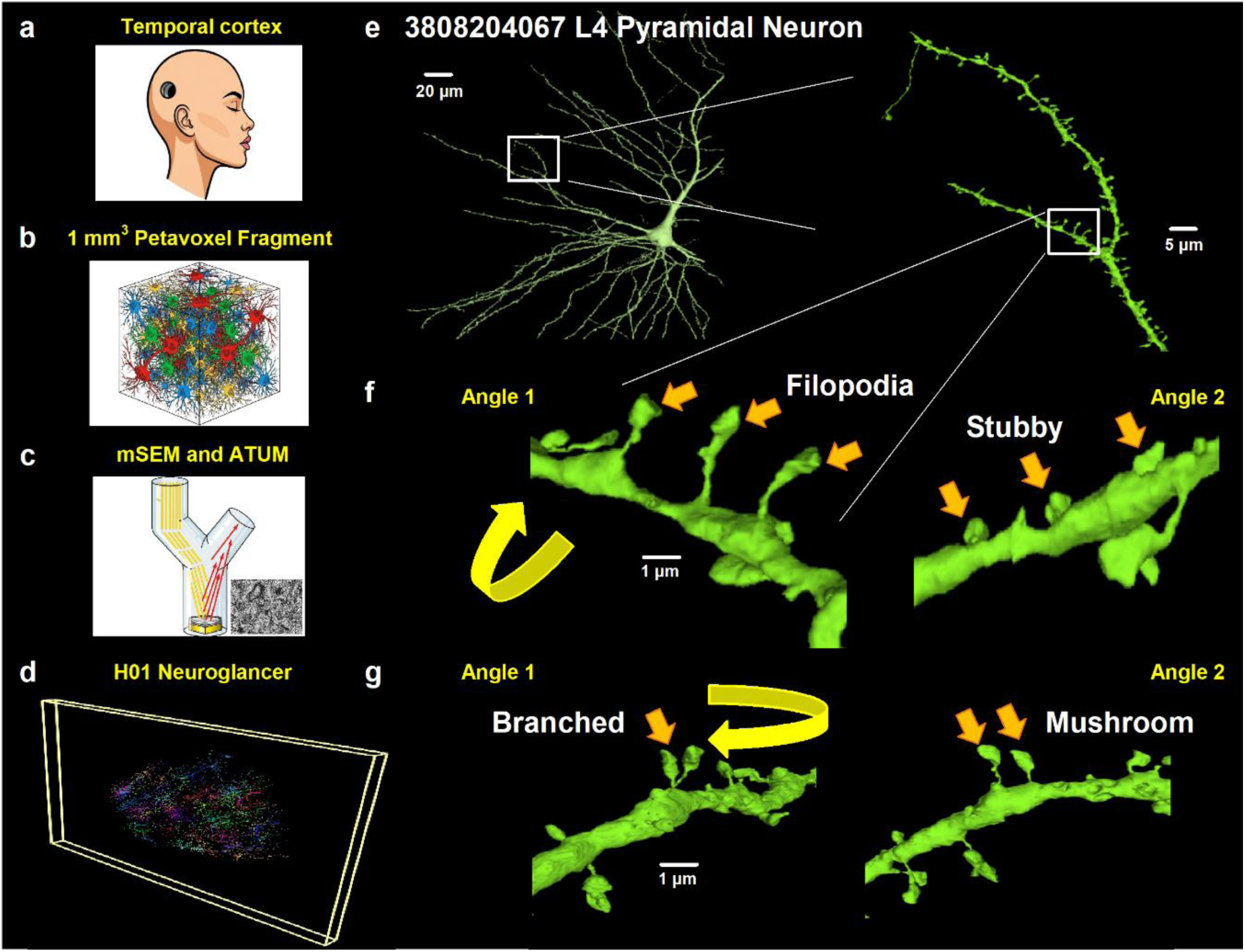
Pipeline from human temporal cortex to three-dimensional spine reclassification, and representative examples of view-dependent classification. (a) Source tissue, a fragment of the anterior middle temporal gyrus of the human cerebral cortex. (b) The reconstructed volume is a cubic millimeter petavoxel fragment containing roughly 57,000 cells and 150 million synapses. (c) Tissue processing and imaging, in which serial ultrathin sections were collected by an Automated Tape-Collecting Ultramicrotome (ATUM) and imaged by multibeam scanning electron microscopy (mSEM). (d) The aligned and segmented H01 volume rendered in the browser-based Neuroglancer viewer. (e) A layer 4 pyramidal neuron, segment ID 3808204067 (Neuroglancer identifier), with two spine-bearing dendritic regions outlined and shown at higher magnification. (f) At Angle 1, three protrusions read as filopodia, and after free rotation to Angle 2, the same three present as neckless stubby spines (orange arrows). (g) A structure scored as a single branched spine at Angle 1 splits into two separate mushroom spines after rotation to Angle 2 (orange arrows). The curved yellow arrow marks the rotation applied between the two views. The schematic illustrations in panels (a), (b), and (c) were created with Mind the Graph. The remaining panels were generated by the authors using Neuroglancer to visualize the H01 dataset (Shapson-Coe et al., 2024).

From a layer 4 pyramidal neuron (Figure 1e, left panel), we retrieved spine-bearing basal dendrites and inspected each protrusion at the zoom level at which it was loaded (Figure 1e, right panel). Figure 1f shows an example of spine-shape classification that depends strongly on viewing angle. The same protrusions that read as filopodia at Angle 1 appeared as neckless, stubby spines when the dendrite was rotated to Angle 2. Curved yellow arrow illustrates the rotation.

Figure 1g illustrates another clear example of spine-shape classification that depends strongly on viewing angle. A structure that appeared as a single-branched spine at Angle 1 split into two separate mushroom spines after rotation to Angle 2. Therefore, rotating nanoscale EM reconstructions of dendrites reveal view-dependent spine-shape classification.

### Quantitative observations

Spine classifications of 445 spines of layer 4 basal dendrites of 5 pyramidal neurons from the two viewing angles were tabulated as a confusion matrix and displayed with a diverging color scale (Figure 2). Each cell was shaded according to its spine count, with blue tones marking concordance, that is, the same designation before and after rotation, and red tones marking discordance, that is, a spine that changed category. Concordant outcomes lie on the diagonal of the matrix, where the Angle 1 and Angle 2 labels coincide, and discordant outcomes lie off the diagonal. The color scale, therefore, makes the result visible at a glance. A faithful taxonomy would concentrate the counts in a deep blue diagonal band, whereas the observed matrix is dominated by red off-diagonal cells.

**Figure 2.**
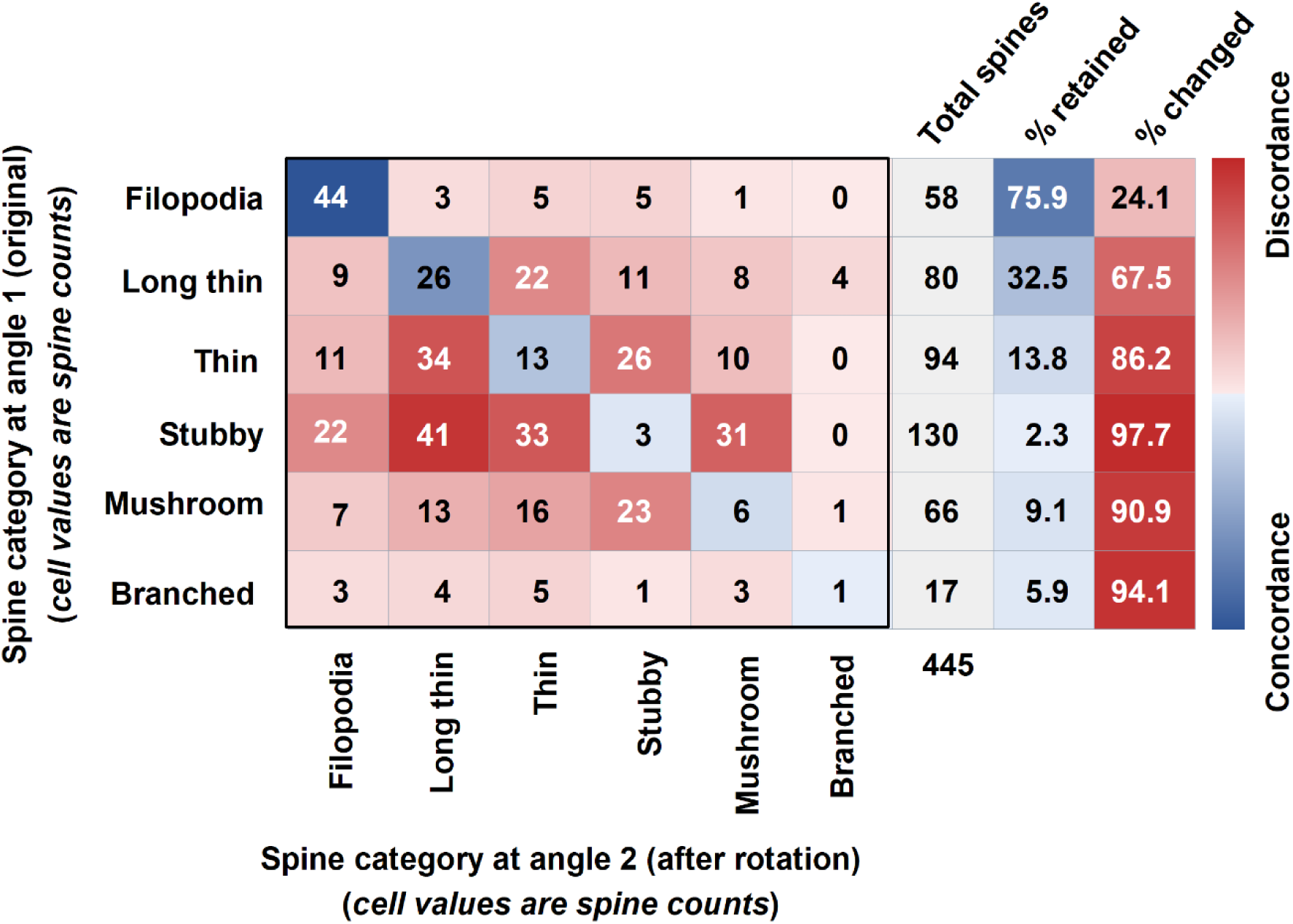
Confusion matrix of dendritic spine classification before (Angle 1, rows) and after (Angle 2, columns) three-dimensional rotation in Neuroglancer. In other words, each spine was classified twice, once from its initial viewpoint (Angle 1) and once after rotation (Angle 2), so the matrix compares two readings of the same dendritic spines. Cells are shaded with a diverging color scale, indicated by the vertical color bar at right, that runs from deep blue for concordance, meaning the same designation before and after rotation, to deep red for discordance, meaning a spine that changed category, with the intensity of each tone scaled to the cell count. Concordant outcomes lie on the diagonal, and discordant outcomes lie off the diagonal. The Total spines column is shaded gray, while the “% retained” and “% changed” columns follow the same diverging scale as the matrix. For each original category, % retained is the diagonal count divided by the row total, and % changed is its complement. See formulas (1) and (2). Concordance was 93 of 445, or 20.9%, and chance-corrected agreement was negligible (Cohen’s kappa = 0.027; Cohen, 1960; Sim and Wright, 2005). The distribution of changes was significantly asymmetric, indicating that reassignment was directional rather than random (McNemar-Bowker test of symmetry, χ² = 54.9, df = 15, p = 1.8 × 10⁻⁶; Bowker, 1948).

In general, cell values are spine counts, but diagonal cells (concordant, blue) indicate spines that retain their category after rotation; off-diagonal cells (discordant, red) indicate spines that are reclassified into a different category. For each original category, “% retained” (illustrated in the right panel of Figure 2) is the diagonal count divided by the row total (e.g., Filopodia: 44/58 = 75.9%), and “% changed” is its complement:

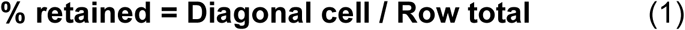

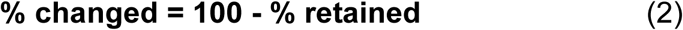

Cell color in Figure 2 encodes the degree of concordance or discordance on the blue-to-red scale, whereas the color of the numbers (white or black) was chosen only for contrast against the cell shade and carries no additional meaning.

Across the L4 dendrites analyzed, a total of 445 paired spine classification events were recorded from the two viewing angles. Only 93 of these events fell on the diagonal of the matrix. The same designation was therefore retained before and after rotation in just 20.9% of cases, while the remaining 79.1% of spines changed category when the dendrite was rotated (Figure 2). Chance-corrected agreement was negligible (Cohen’s kappa = 0.027). The level of stable classification was thus barely above what would be expected if labels had been assigned at random.

Instability was not shared equally across the six categories. For each original category, the fraction of spines that retained their designation after rotation and the fraction that changed are reported alongside the matrix (Figure 2, right). Filopodia were the most stable category, with 75.9% retained and 24.1% changed, which is consistent with their distinctive elongated form being recognizable from most viewpoints.

All conventional head-bearing classes were far less stable. Long-thin spines retained their designation in 32.5% of cases and changed in 67.5%. Thin spines were retained in 13.8% and changed in 86.2%. Mushroom spines were retained in only 9.1% and changed in 90.9%. Branched spines were retained at 5.9% and changed in 94.1%. Stubby spines were the least stable of all, with just 2.3% retained and 97.7% changed. The classes on which most functional interpretation rests are therefore precisely the ones whose assignment is least reproducible across viewing angles.

The changes in designation were not random. A McNemar-Bowker test of the confusion matrix rejected marginal symmetry (χ2 = 54.9, df = 15, p = 1.8 x 10^-6^). Reassignment, therefore, occurred preferentially along particular axes of the table, and the densest red cells identify these preferred transitions. Categories sharing similar head-and-neck geometry were the most readily interchanged. Spines initially scored as stubby were frequently reassigned after rotation to long thin (41 events), thin (33 events), or mushroom (31 events). This is consistent with the expectation that a foreshortened thin or mushroom spine viewed end on mimics a neckless stubby protrusion.

Likewise, spines scored as thin at Angle 1 commonly shifted to long thin (34 events) at Angle 2, indicating that these two categories share morphological features that are difficult to distinguish from a single perspective. The marginal totals shifted accordingly. Stubby was the most frequent class at Angle 1, with 130 events, but dropped sharply at Angle 2, with 69 events. Long thin rose from 80 to 121 events. The apparent prevalence of stubby spines is therefore in part an artifact of viewing angle.

Taken together, the matrix statistics and the rendered examples show that the conventional classification of human cortical spines depends strongly and systematically on the angle of observation, with four of every five spines changing category under rotation alone.

## Discussion

This study used dendrite rotation in a nanoscale EM reconstruction to isolate the effect of viewing angle on spine-shape classification. The central finding is unambiguous. Only about one spine in five retained its conventional category when its parent dendrite was rotated, chance-corrected agreement was negligible, and the reclassifications were systematic rather than random. Stubby and thin spines were frequently revealed as mushroom or long thin spines once viewed from a second angle. Because the object, the observer, and the criteria were held constant, these changes can be attributed solely to viewpoint.

The importance of this result lies in what it reveals about the technique on which most spine classifications still depend. Golgi-Cox impregnation has genuine and well-known strengths. It is inexpensive, it applies to virtually any tissue, including post-mortem human brain, and it yields long-lived preparations (Risher et al., 2014; Flores et al., 2025). However, it also has an intrinsic and rarely acknowledged weakness. The orientation of a spine in the histological section is set by the accident of how its dendrite lies in the slice, and it cannot be controlled by the experimenter. The same is true for confocal and two-photon imaging, with the added penalty of poor axial resolution (Pchitskaya and Bezprozvanny, 2020).

It is important to clarify how our approach differs from previous EM studies of spine morphology. In those studies, individual spines were reconstructed and examined in three dimensions to recover their true geometry, head volume, surface area, neck length, and neck diameter, with rotation serving to remove the viewing angle and measure each spine free of projection (Arellano et al., 2007; Ofer et al., 2021). Our design inverts this purpose. We rotated the parent dendrite together with its spines, not each spine in isolation, and we did so not to measure intrinsic geometry but to preserve and expose the dependence of classification on viewpoint. The unit of analysis is therefore the dendrite as the observer encounters it, rather than the isolated spine, and the rotation varies the viewing angle instead of eliminating it. This lets us ask a question those studies did not pose: whether the category assigned to a spine is stable when the same object is seen from a second orientation. To our knowledge, no prior EM study has rotated intact spine-bearing dendrites to test the reproducibility of conventional shape classification, which makes the present analysis distinct in both its design and its question from earlier three-dimensional reconstructions of spines.

We did not analyze Golgi-Cox or confocal preparations directly. Instead, we emulated their single-viewpoint geometry by classifying each spine using a single arbitrary orientation of the rotatable H01 petavoxel fragment. Because both techniques classify dendritic spines from a single uncontrolled projection, the instability we measured under these emulated conditions is inherent to their design rather than specific to our dataset.

Methods designed to make Golgi-Cox classification faster and more objective, such as the geometry-based scheme of Risher et al. (2014), standardize measurement procedures. However, they also cannot remove the orientation problem because they still operate on a single, uncontrolled projection. The present data show that this is not a marginal source of noise. It flips most classifications and, at the population level, it inflates the apparent frequency of stubby spines while reducing that of mushroom and long-thin spines. Quantitative claims about the proportions of spine types derived from single-view preparations should therefore be treated with caution.

Although considerable effort has been devoted to classifying dendritic spines in human tissue (Ofer et al., 2022; Renner and Rasia-Filho, 2023; Schünemann et al., 2025), most of these approaches rely on confocal microscopy for image acquisition and reconstruction. Schünemann et al. (2025) classified nearly 4,000 human dendritic spines into mushroom, thin, and stubby subtypes using confocal Z-stacks combined with deep learning-aided reconstruction, demonstrating the feasibility of large-scale analysis with this modality. However, the authors acknowledge that confocal microscopy imposes important resolution constraints. Because spine heads frequently measure below 1,000 nm while confocal systems resolve only around 200 to 300 nm, smaller spines, and particularly the slender neck region critical to most classification schemes, may be obscured or missed. Their own data reflect this, as the spine neck was the structure most often incompletely captured, even by their trained algorithm.

In contrast, our methodology provides higher resolution and overcomes several of these limitations. The improved resolution allows reliable detection of the spine, neck, and head, the structures that confocal imaging tends to underrepresent, thereby reducing the systematic loss of small or thin spines that biases density and subtype distributions. Furthermore, confocal Z-stacks capture spines in a fixed orientation and cannot be freely rotated, so a spine may be measured from a single viewing angle that does not reflect its true dimensions. This constraint can distort head and neck measurements and, because classification depends on ratios derived from these values, introduce bias into the assignment of morphological spine subtypes. Our approach permits full dendrite rotation and inspection of each spine in three dimensions, enabling measurements that are independent of viewing angle and yielding a more accurate and unbiased classification of human dendritic spine morphology.

Recent computational efforts point in the same direction. Smirnova et al. (2026) moved beyond scalar size metrics by introducing three-dimensional shape descriptors based on spherical harmonics and Zernike moments and used unsupervised clustering rather than fixed labels to separate control from Alzheimer model spines. Their descriptors capture morphology more sensitively than the named classes do, and they treat spine shape as a continuum by modeling strategies derived from analyses of rigid confocal images. Yet their meshes are still reconstructed from confocal Z stacks, and the spherical harmonic representation is not intrinsically rotation invariant, so each spine must be computationally realigned to a common orientation before it can be compared. However, the approach taken in our study is more direct. Because we analyze every spine as an explicit nanometer-scale mesh in the H01 reconstruction, which can be rotated freely in Neuroglancer, we solve the orientation problem with the data themselves rather than with a downstream alignment-modeling algorithm.

The originality of our approach is that it provides direct within-object evidence of a continuum, thereby complementing the distributional evidence already available. Earlier three-dimensional studies inferred a continuum from the unimodal shape of population distributions of head volume and neck dimensions in mouse and human cortex (Ofer et al., 2021, 2022), and from the inability of segmentation pipelines to recover separable clusters (Pchitskaya and Bezprozvanny, 2020). Those analyses argue convincingly that discrete classes are not visible in aggregate statistics.

The dendritic rotational analysis presented here makes a logically distinct and more pointed argument. It shows that the assignment of an individual spine to a class is itself unstable, because the diagnostic features, namely head width, neck length, and the length-to-width ratio, are projection-dependent quantities. A taxonomy whose categories interconvert under rotation cannot describe intrinsic kinds. This is the strongest sense in which previous Golgi-Cox classifications can be said to be wrong. It is not that the measurements were carelessly made, but that the entities they purported to measure are partly artifacts of the viewpoint.

Several limitations should be noted. The analysis was performed on a single human specimen that was resected for epilepsy, and although it was reported as histologically normal, subtle effects of the disease or its treatment cannot be excluded (Shapson-Coe et al., 2024). Classification at both angles was carried out by a trained observer, so variability between observers, which is itself a well-documented problem for Golgi-Cox analysis, was not assessed here and would be expected to add further to the disagreement. The two-angle design quantifies the minimum instability produced by a single rotation, and sampling many orientations per spine would likely reveal even greater dependence on viewing angle. Finally, the present report is confined to demonstrating bias by emulating the uncontrolled viewing geometry of conventional Golgi-Cox or confocal preparations; however, additional studies are necessary to measure changes in the length, area, and volume of the identified spines after dendrite rotation.

Neuroglancer exposes the full three-dimensional mesh and voxel data for each spine. Intrinsically orientation-independent quantities, such as head and neck volumes, surface areas, sphericity, and the geometry of the neck, can therefore be extracted, and the spines can be clustered in a continuous feature space without recourse to named shapes. This approach has already been advocated on theoretical grounds (Pchitskaya and Bezprozvanny, 2020) but not using the EM H01 fragment (Shapson-Coe et al., 2024). Our study proposes that recasting spine morphology as a measurable continuum, anchored in rotation-invariant three-dimensional features rather than in viewpoint-dependent silhouettes, offers a path toward classifications that are reproducible across observers, laboratories, and imaging modalities. Such classifications can also be more faithfully related to the study of synaptic function in health and disease.

## Conclusions

A single spine viewed from a single angle is not a spine class or category. It is a shadow of one. By rotating the same human cortical dendrites containing spines in three dimensions, this study shows that four of every five spines change category with viewpoint alone, and that the changes are directional rather than random. The discrete labels that have organized a century of work, from Cajal’s first drawings to modern Golgi-Cox and confocal studies, therefore confound the true shape of a spine with the arbitrary angle from which it is viewed. Nanoscale EM reconstructions, such as H01 and Neuroglancer, allow every dendrite and its spines to be turned freely and measured by quantities that do not depend on the observer. The future of spine morphology, therefore, lies not in deciding which of six names a protrusion deserves, but in placing it within a continuous, rotation-invariant space of shape. These reconstructions, explored in tools like Neuroglancer, will reveal precisely how spine morphology is modified during learning and memory, as well as in pathological processes.

## Author contributions

EM conceived and designed the study and wrote the paper. All authors analyzed the database, revised the manuscript, and approved it.

## Ethical Approval Statement

This manuscript does not require an ethical approval statement, as it was performed on a publicly available database.

## References

Aguilar-Hernández L, Flores-Gómez GD, Nacher J, Morales-Medina JC, Flores G. Cerebrolysin Ameliorates Age-Induced Dendritic Spine Degeneration and Memory Decline in C57BL6 Mice. Neurochem Res. 2025 Dec 29;51(1):23. doi: 10.1007/s11064-025-04627-0. PMID: 41460391; PMCID: PMC12748144.

Arellano, J. I., Benavides-Piccione, R., Defelipe, J., & Yuste, R. (2007). Ultrastructure of dendritic spines: correlation between synaptic and spine morphologies. Frontiers in neuroscience, 1(1), 131–143. 10.3389/neuro.01.1.1.010.2007

Benavides-Piccione R, Fernaud-Espinosa I, Robles V, Yuste R, DeFelipe J. Age-based comparison of human dendritic spine structure using complete three-dimensional reconstructions. Cereb Cortex. 2013 Aug;23(8):1798–810. doi: 10.1093/cercor/bhs154. Epub 2012 Jun 17. PMID: 22710613; PMCID: PMC3698364.

Berry, K. P., & Nedivi, E. (2017). Spine Dynamics: Are They All the Same?. Neuron, 96(1), 43–55. 10.1016/j.neuron.2017.08.008

Bhatt, D. H., Zhang, S., & Gan, W. B. (2009). Dendritic spine dynamics. Annual review of physiology, 71, 261–282. 10.1146/annurev.physiol.010908.163140

Bokota, G., Magnowska, M., Kuśmierczyk, T., Łukasik, M., Roszkowska, M., & Plewczynski, D. (2016). Computational Approach to Dendritic Spine Taxonomy and Shape Transition Analysis. Frontiers in computational neuroscience, 10, 140. 10.3389/fncom.2016.00140

Bourne, J. N., & Harris, K. M. (2008). Balancing structure and function at hippocampal dendritic spines. Annual review of neuroscience, 31, 47–67. 10.1146/annurev.neuro.31.060407.125646

Bowker AH (1948). A test for symmetry in contingency tables. Journal of the American Statistical Association, 43(244), 572–574. 10.1080/01621459.1948.10483284

Cabrera-Pedraza FJ, Apam-Castillejos DJ, de la Cruz-López F, Garcés-Ramírez L, Tendilla-Beltrán H, Flores G. Quetiapine treatment corrects behavioral impairments and promotes neuroplasticity in a neurodevelopmental rat model of schizophrenia. Neuroscience. 2026 Jan 9;592:223–231. doi: 10.1016/j.neuroscience.2025.11.036. Epub 2025 Nov 28. PMID: 41319804.

Choi, J., Lee, S. E., Lee, Y., Cho, E., Chang, S., & Jeong, W. K. (2023). DXplorer: A Unified Visualization Framework for Interactive Dendritic Spine Analysis Using 3D Morphological Features. IEEE transactions on visualization and computer graphics, 29(2), 1424–1437. 10.1109/TVCG.2021.3116656

Cohen J (1960) A coefficient of agreement for nominal scales. Educ Psychol Meas 20:37–46.

Dai J, Zeng Q, Cheng L, Chen H, Jiang L, Hu Y. Biphasic response of silent synapses: Differential remodeling of hippocampal synaptic plasticity by status epilepticus in juvenile versus adult mice. Neurochem Int. 2026 May;195:106147. doi: 10.1016/j.neuint.2026.106147. Epub 2026 Mar 11. PMID: 41825802.

Escribano-Cadena Y, Tendilla-Beltrán H, Flores G. Alterations in dendritic spine plasticity in the prefrontal cortex induced by social play restrictions in male rats. Neuroscience. 2025 Sep 13;583:23–32. doi: 10.1016/j.neuroscience.2025.07.041. Epub 2025 Jul 27. PMID: 40730251.

Ferreira A, Constantinescu VS, Malvaut S, Saghatelyan A, Hardy SV. Distinct forms of structural plasticity of adult-born interneuron spines in the mouse olfactory bulb induced by different odor learning paradigms. Commun Biol. 2024 Apr 6;7(1):420. doi: 10.1038/s42003-024-06115-7. PMID: 38582915; PMCID: PMC10998910.

Flores G, Aguilar-Hernández L, García-Dolores F, Nicolini H, Vázquez-Hernández AJ, Tendilla-Beltrán H. Dendritic spine degeneration: a primary mechanism in the aging process. Neural Regen Res. 2025 Jun 1;20(6):1696–1698. doi: 10.4103/NRR.NRR-D-24-00311. Epub 2024 Jun 26. PMID: 39104099; PMCID: PMC11688554.

García-López, P., García-Marín, V., & Freire, M. (2010). Dendritic spines and development: towards a unifying model of spinogenesis--a present day review of Cajal’s histological slides and drawings. Neural plasticity, 2010, 769207. 10.1155/2010/769207

Gilles, J. F., Mailly, P., Ferreira, T., Boudier, T., & Heck, N. (2024). Spot Spine, a freely available ImageJ plugin for 3D detection and morphological analysis of dendritic spines. F1000Research, 13, 176. 10.12688/f1000research.146327.2

Hu, X., Guo, Z., Shi, Z., Zhen, P., & Zhou, M. (2025). Morphological changes in CA3 pyramidal neurons after transient global ischemia. Neuroreport, 36(14), 856–863. 10.1097/WNR.0000000000002206

Kashiwagi Y, Higashi T, Obashi K, et al.: Computational geometry analysis of dendritic spines by structured illumination microscopy. Nat. Commun. 2019;10(1):1285. 10.1038/s41467-019-09337-0

Kayedi-Bakhtiari N, Vatanparast J. Early handling improves sociability and preference for social novelty in social-defeated male rats: Laterality in vermal Purkinje cells architecture. Behav Brain Res. 2025 Oct 2;494:115714. doi: 10.1016/j.bbr.2025.115714. Epub 2025 Jul 4. PMID: 40618962.

Loewenstein, Y., Kuras, A., & Rumpel, S. (2011). Multiplicative dynamics underlie the emergence of the log-normal distribution of spine sizes in the neocortex in vivo. The Journal of neuroscience: the official journal of the Society for Neuroscience, 31(26), 9481–9488. 10.1523/JNEUROSCI.6130-10.2011

Luengo-Sanchez S, Fernaud-Espinosa I, Bielza C, Benavides-Piccione R, Larrañaga P, DeFelipe J. 3D morphology-based clustering and simulation of human pyramidal cell dendritic spines. PLoS Comput Biol. 2018 Jun 13;14(6):e1006221. doi: 10.1371/journal.pcbi.1006221. PMID: 29897896; PMCID: PMC6060563.

Masoli S, Rizza MF, Moccia F, D’Angelo E. The spiny relationship between parallel fibers, climbing fibers, and Purkinje cells. Front Physiol. 2025 Oct 9;16:1671271. doi: 10.3389/fphys.2025.1671271. PMID: 41141850; PMCID: PMC12546079.

Ofer, N., Berger, D. R., Kasthuri, N., Lichtman, J. W., & Yuste, R. (2021). Ultrastructural analysis of dendritic spine necks reveals a continuum of spine morphologies. Developmental neurobiology, 81(5), 746–757. 10.1002/dneu.22829

Ofer, N., Benavides-Piccione, R., DeFelipe, J., & Yuste, R. (2022). Structural Analysis of Human and Mouse Dendritic Spines Reveals a Morphological Continuum and Differences across Ages and Species. eNeuro, 9(3), ENEURO.0039-22.2022. 10.1523/ENEURO.0039-22.2022

Otero PA, Fricklas G, Nigam A, Lizama BN, Wills ZP, Johnson JW, Chu CT. Endogenous PTEN-Induced Kinase 1 Regulates Dendritic Architecture and Spinogenesis. J Neurosci. 2022 Oct 12;42(41):7848–7860. doi: 10.1523/JNEUROSCI.0785-22.2022. Epub 2022 Sep 7. PMID: 36414008; PMCID: PMC9581559.

Paul S, Pramanick R, Das N, Baczynska E, Bedrood Z, Chakraborti T, Basu S, Wlodarczyk J. 2dSpAn-Auto: an automated tool for analysis of two-dimensional dendritic spine images. BMC Bioinformatics. 2025 Jul 1;26(1):162. doi: 10.1186/s12859-025-06179-0. PMID: 40597607; PMCID: PMC12211165.

Peters, A., & Kaiserman-Abramof, I. R. (1970). The small pyramidal neuron of the rat cerebral cortex. The perikaryon, dendrites and spines. The American journal of anatomy, 127(4), 321–355. 10.1002/aja.1001270402

Pchitskaya, E., & Bezprozvanny, I. (2020). Dendritic Spines Shape Analysis-Classification or Clusterization? Perspective. Frontiers in synaptic neuroscience, 12, 31. 10.3389/fnsyn.2020.00031

Pchitskaya, E., Vasiliev, P., Smirnova, D., Chukanov, V., & Bezprozvanny, I. (2023). SpineTool is an open-source software for analysis of morphology of dendritic spines. Scientific reports, 13(1), 10561. 10.1038/s41598-023-37406-4

Portera-Cailliau C, Pan DT, Yuste R. Activity-regulated dynamic behavior of early dendritic protrusions: evidence for different types of dendritic filopodia. J Neurosci. 2003 Aug 6;23(18):7129–42. doi: 10.1523/JNEUROSCI.23-18-07129.2003. PMID: 12904473; PMCID: PMC6740658.

Reberger, R., Dall’Oglio, A., Jung, C. R., & Rasia-Filho, A. A. (2018). Structure and diversity of human dendritic spines evidenced by a new three-dimensional reconstruction procedure for Golgi staining and light microscopy. Journal of neuroscience methods, 293, 27–36. 10.1016/j.jneumeth.2017.09.001

Reimer, M. L., Kauer, S. D., Benson, C. A., King, J. F., Patwa, S., Feng, S., Estacion, M. A., Bangalore, L., Waxman, S. G., & Tan, A. M. (2024). A FAIR, open-source virtual reality platform for dendritic spine analysis. Patterns (New York, N.Y.), 5(9), 101041. 10.1016/j.patter.2024.101041

Renner, J., & Rasia-Filho, A. A. (2023). Morphological Features of Human Dendritic Spines. Advances in neurobiology, 34, 367–496. 10.1007/978-3-031-36159-3_9

Risher, W. C., Ustunkaya, T., Singh Alvarado, J., & Eroglu, C. (2014). Rapid Golgi analysis method for efficient and unbiased classification of dendritic spines. PloS one, 9(9), e107591. 10.1371/journal.pone.0107591

Rochefort, N. L., & Konnerth, A. (2012). Dendritic spines: from structure to in vivo function. EMBO reports, 13(8), 699–708. 10.1038/embor.2012.102

Rodriguez A, Ehlenberger DB, Dickstein DL et al. Automated three-dimensional detection and shape classification of dendritic spines from fluorescence microscopy images. PLoS One 2008;3:e1997.

Schünemann, K. D., Hattingh, R. M., Verhoog, M. B., Yang, D., Bak, A. V., Peter, S., van Loo, K. M. J., Wolking, S., Kronenberg-Versteeg, D., Weber, Y., Schwarz, N., Raimondo, J. V., Melvill, R., Tromp, S. A., Butler, J. T., Höllig, A., Delev, D., Wuttke, T. V., Kampa, B. M., & Koch, H. (2025). Comprehensive analysis of human dendritic spine morphology and density. Journal of neurophysiology, 133(4), 1086–1102. 10.1152/jn.00622.2024

Shapson-Coe A, Januszewski M, Berger DR, Pope A, Wu Y, Blakely T, Schalek RL, Li PH, Wang S, Maitin-Shepard J, Karlupia N, Dorkenwald S, Sjostedt E, Leavitt L, Lee D, Troidl J, Collman F, Bailey L, Fitzmaurice A, Kar R, Field B, Wu H, Wagner-Carena J, Aley D, Lau J, Lin Z, Wei D, Pfister H, Peleg A, Jain V, Lichtman JW. (2024). A petavoxel fragment of human cerebral cortex reconstructed at nanoscale resolution. Science (New York, N.Y.), 384(6696), eadk4858. 10.1126/science.adk4858

Sim J, Wright CC. The kappa statistic in reliability studies: use, interpretation, and sample size requirements. Phys Ther. 2005 Mar;85(3):257–68. PMID: 15733050.

Smirnova, D., Ustinova, A., Chukanov, V., & Pchitskaya, E. (2026). 3D dendritic spines shape descriptors for efficient classification and morphology analysis in control and Alzheimer’s disease modeling neurons. Bioinformatics (Oxford, England), 42(2), btag025. 10.1093/bioinformatics/btag025

Tendilla-Beltrán H, Antonio Vázquez-Roque R, Judith Vázquez-Hernández A, Garcés-Ramírez L, Flores G. Exploring the Dendritic Spine Pathology in a Schizophrenia-related Neurodevelopmental Animal Model. Neuroscience. 2019 Jan 1;396:36–45. doi: 10.1016/j.neuroscience.2018.11.006. Epub 2018 Nov 16. PMID: 30452973.

Tennin M, Matkins HT, Rexrode L, Bollavarapu R, Pareek T, Kroeger D, Pantazopoulos H, Gisabella B. Sleep deprivation alters hippocampal dendritic spines in a contextual fear memory engram. Sci Rep. 2026 Feb 24;16(1):10381. doi: 10.1038/s41598-026-41336-2. PMID: 41735509; PMCID: PMC13031319.

Theobald CC, Lotfinia A, Knobloch JA, Medlej Y, Stevens DR, Lauterbach MA. Distribution of spine classes shows intra-neuronal dendritic heterogeneity in mouse cortex. Neurophotonics. 2025 Jan;12(1):015001. doi: 10.1117/1.NPh.12.1.015001. Epub 2024 Dec 19. PMID: 39712647; PMCID: PMC11657875.

Tønnesen, J., & Nägerl, U. V. (2016). Dendritic Spines as Tunable Regulators of Synaptic Signals. Frontiers in psychiatry, 7, 101. 10.3389/fpsyt.2016.00101

Urban, P., Rezaei, V., Bokota, G., Denkiewicz, M., Basu, S., & Plewczyński, D. (2019). Dendritic Spines Taxonomy: The Functional and Structural Classification • Time-Dependent Probabilistic Model of Neuronal Activation. Journal of computational biology: a journal of computational molecular cell biology, 26(4), 322–335. 10.1089/cmb.2018.0155

Ustinova A, Volkova E, Rakovskaya A, Smirnova D, Korovina O, Pchitskaya E. Generate and Analyze Three-Dimensional Dendritic Spine Morphology Datasets With SpineTool Software. Curr Protoc. 2024 Dec;4(12):e70061. doi: 10.1002/cpz1.70061. PMID: 39641661.

Wilson, C. J., Groves, P. M., Kitai, S. T., & Linder, J. C. (1983). Three-dimensional structure of dendritic spines in the rat neostriatum. The Journal of neuroscience: the official journal of the Society for Neuroscience, 3(2), 383–388. 10.1523/JNEUROSCI.03-02-00383.1983

Yuste R. The discovery of dendritic spines by Cajal. Front Neuroanat. 2015 Apr 21;9:18. doi: 10.3389/fnana.2015.00018. PMID: 25954162; PMCID: PMC4404913.

